# Adenine nucleotide translocase 2 silencing promotes metabolic adaptations and anoikis in P19 embryonal carcinoma stem cells

**DOI:** 10.1101/2024.06.07.597817

**Authors:** Gabriela L. Oliveira, Sandra I. Mota, Paulo J. Oliveira, Ricardo Marques

## Abstract

Cancer stem cells (CSCs) are amongst the group of cells constituting tumors, being characterized by their strong self-renewal and survival properties. Cancer cells, CSCs included, are thought to rely mostly on glycolysis, even in the presence of oxygen, which confers them adaptative advantages. Adenine nucleotide translocator 2 (ANT2), responsible for the exchange of ADP and ATP in the mitochondrial inner membrane, has been correlated with a higher glycolytic metabolism and is known to be overexpressed in cancer cells. Using P19 embryonal carcinoma stem cells (P19SCs) as a CSCs model, we inhibit ANT2 translation by using siRNA. ANT2 protein levels were shown to be overexpressed in P19SC when compared to their differentiated counterparts. Furthermore, we showed here that the OXPHOS machinery and mitochondrial membrane potential are compromised after ANT2 depletion, exhibiting a metabolic adaptation towards a less oxidative phenotype. Interestingly, hexokinase II levels were downregulated, which was also accompanied by decreased cell growth, and decreased ability to form spheroids. Our findings underscore ANT2 as a key regulator of metabolic remodeling and cell survival of CSCs, suggesting its potential as a therapeutic target for controlling CSC-driven tumor progression.

**Highlights:**

- ANT2 silencing promotes cell growth arrest and metabolic remodeling in CSCs.
- ANT2 depletion modulates HKII protein levels.
- ANT2 induce anoikis resistance in P19SCs

## 1. Introduction

Cancer is a multifactorial and life-threatening condition in which poor outcomes are related to the metastatic potential of tumors, despite the advances in available treatments. A hallmark of cancer is metabolic reprogramming, which confers adaptive advantages to cancer cells by enabling proliferation and survival even under adverse conditions, thus facilitating tumor spread.

In recent years, cancer stem cells (CSCs) have emerged as architects and building blocks of the survival and progression of various tumor types. This niche population possesses self-renewal capacity, tumorigenic potential, and high plasticity due to the ability to differentiate into specialized cell types that promote tumor heterogeneity, which ultimately is thought to drive tumor recurrence, relapse, and metastasis [1]. Despite advances in cancer treatments, conventional radio- and chemotherapies, often fail to effectively target these elusive cells, thus ultimately contributing to high mortality rates [2–4]. A deeper understanding of CSCs’ biological characteristics and molecular mechanisms holds promise for the development of novel therapeutics [5].

P19 embryonal carcinoma stem cells (P19SCs) are a suitable model to study stem cell maintenance and differentiation [6,7], and also a cancer stem cell-like model for CSCs, providing insights into therapy evasion, metabolic phenotype, or mitochondrial adaptations [8,9]. Emerging research underlines the distinctive metabolic behavior of CSCs, as they display an increased dependence on glycolysis in contrast to their differentiated counterparts. This altered metabolic preference enables CSCs to thrive in nutrient-deprived environments and withstand therapeutic stress [2,3]. In our previous works, we demonstrated that P19SCs have a glycolysis-dependent phenotype while differentiation promotes a metabolic shift towards mitochondrial metabolism and mitochondrial remodeling, which may also be accompanied by changes in the adenine nucleotide translocator (ANT) expression levels [8].

ANT, a nuclear-encoded integral protein located in the inner mitochondrial membrane, imports ADP into the mitochondrial matrix, which in turn is used to generate ATP by oxidative phosphorylation (OXPHOS) under physiological conditions. ATP can then be dispatched from the mitochondria into the cytosol, where it can be utilized for the metabolic needs of the cell [10]. There are four known isoforms of ANT in humans (ANT1-4), with expression depending on the cell type, tissue nature, developmental stage, cell proliferation status, as well as energy requirements [11]. ANT1 is primarily found in differentiated tissues, such as skeletal muscle, heart, and brain [12]. ANT2 is preferentially expressed in undifferentiated cells or regenerative tissues, such as the kidney and liver [13]. ANT3 is ubiquitously expressed, and ANT4 is exclusively localized in germ cells [14]. Additionally, ANT plays a role in cell death pathways by regulating mitochondrial membrane permeability and cell fate decisions, with some isoforms acting as pro-apoptotic (ANT1 and 3) while others function as anti-apoptotic (ANT2 and 4) [15]. It has been suggested that under mitochondrial respiration impairment, ANT2 imports ATP produced by glycolysis into the mitochondria, where it is used by the F1Fo-ATPase complex to eject protons into the intermembrane space, in contrast to what happens under physiological conditions [16]. This process enables the maintenance of mitochondrial membrane potential (ΔΨm) and ensures cell survival from induced apoptosis [17]. Therefore, ANT2 expression is associated with growth-dependence, is considered a cell proliferation marker [18], and plays a role in cancer. Indeed, evidence suggests that ANT2 is essential during tumor development which may be related to metabolic adaptation, and linked to glycolytic metabolism [13,19]. Accordingly, it is highly upregulated in various cancer cells and hormone-dependent cancers [20], and its silencing increases apoptosis and decreases cell growth in human breast cancer, and breast cancer stem-like cells [21,22]. Despite the evidence, little is known regarding the presence and functional effects of ANT2 in cancer cells expressing a stem cell-like phenotype, its impacts on cell metabolism, and how it influences mitochondrial activity.

The current work investigated the presence and functional role of ANT2 in the intrinsic capabilities of P19SCs by assessing stem cell growth, metastasis, therapy evasion, and metabolic effects on cancer progression. Based on our findings, which are supported by mounting evidence, ANT2 emerges as a pivotal element for mitochondrial function, resistance to cell detachment-induced apoptosis (anoikis), and metabolic adaptations of CSCs, making it a promising target for therapeutic intervention.

## 2. Material and Methods

### 2.1 Cell culture and differentiation

P19 embryonal carcinoma cell line was obtained from the American Type Culture Collection (ATCC® CRL-1825™, RRID:CVCL_2153) and cultured in High glucose Dulbecco’s modified Eagle’s medium (DMEM-D5648, Sigma-Aldrich) supplemented with 10% FBS, 1.8 g/l sodium bicarbonate, 110 mg/l sodium pyruvate, and 1% antibiotic/antimycotic solution, at 37°C in a humidified atmosphere of 5% CO_2_. Cells were passaged every 2-3 days when reaching 70-80% confluence, at a 1:20 to 1:30 dilution. To induce cell differentiation, P19 cells were seeded at a density of 5.2x10^3^ cells/cm^2^ and 1 μM of Retinoic Acid was added for 96h, as described previously [6,8].

### 2.2 Cell transfection

Cells were seeded at a density of 7.5x10^3^ cells/cm^2^ and transfected after 24h with 50 nM ANT2 siRNA (siANT2, Mm_Slc25a5_1), sense strand 5’ CGA GCU GCC UAC UUU GCU ATT 3’, or with scrambled siRNA with sense strand 5’ UUC UCC GAA CGU GUC ACG UdT dT 3’, without homology, as negative control (siCtrl), both purchased from Qiagen. Lipofectamine 2000 was used as a transfection agent, according to the manufacturer’s instructions. Briefly, cells were incubated with the siRNA-lipid complex for 5 hours, after which the medium was replaced with fresh growth medium. Transfection efficiency and ANT2 expression were assessed by Western blot analysis at several timepoints post-transfection.

### 2.3 Cell viability/proliferation

P19SCs were plated on 96 multi-well plates with a cellular density of 7.5x10^3^ cells/cm^2^ and transfected after 24 hours with scrambled and ANT2 siRNA, as previously described. After 24, 48, 72, and 96 hours of transfection, fresh medium with 10 μg/ml of resazurin was added and incubated for 90 min in the dark at 37°C a 5% CO_2_ atmosphere. The amount of fluorescence (resorufin) was measured at 540/590 nm, using Cytation3^TM^ UV–vis multi-well plate imaging reader (BioTek) [23]. The plate was washed with PBS 1X and cells were fixed with ice-cold 1% (v/v) acetic acid in methanol overnight at -20°C. Sulforhodamine B (SRB) solution (0,5% in 1% acetic acid) was added to dry wells and incubated at 37°C for 1 hour. After the removal of SRB excess with 1% (v/v) acetic acid, the dye was resuspended in a 10 mM Tris pH 10 solution. The absorbance was then measured at 510 nm and 620 nm, using Cytation3^TM^ UV–vis multi-well plate imaging reader [24].

### 2.4 Western Blot

After harvesting by trypsinization and washing with cold PBS 1X, the cell pellet was resuspended in RIPA buffer supplemented with 2mM DTT, 100 mM PMSF, 20mM NAF, 80 mM Na_3_VO_4_ and protease inhibitor cocktail (1:100), physically disrupted by sonication, and kept at - 80°C until used. The protein content was measured using BCA assay. Twenty micrograms of protein were loaded in a 10 or 12% SDS-polyacrylamide gel and separated by electrophoresis at 120V. Samples were electrotransferred to a polyvinylidene difluoride (Bio-Rad, 1704273) or nitrocellulose (Bio-Rad, 1704271) membrane at 25V, for 10 min, using the Trans-Blot® Turbo^TM^ Transfer System (Bio-Rad Laboratories). Membranes were stained with Ponceau S staining to ensure equal loading and quantification purposes for P19dCs given the observed changes in metabolic- and cytoskeleton-related housekeeping proteins [6]. After blocking the membranes with 5% non-fat dry milk or BSA in TBS-T (50 mM Tris-HCl, pH 7.5, 150 mM NaCl, and 0.1% Tween-20) for 1 h at room temperature, membranes were incubated at 4°C overnight with the primary antibodies (Table 1). After three 10-minute washes with TBS-T, the membranes were incubated with the respective horseradish peroxidase-conjugated secondary antibody (1:5000) for 1 hour at room temperature. Following washes with TBS-T, membranes were incubated with Clarity™ Western ECL Substrate detection kit, and images were acquired in a VWR Gel Documentation System Imager Chemi 5QE. Band densities were determined using TotalLab TL120 software.

**Table 1.** Antibodies list.

| Antibody | Reference |
| --- | --- |
| Alexa Fluor-594 | A11005 Invitrogen. RRID:AB_141372 |
| Alexa Fluor-488 | A11008 Invitrogen. RRID:AB_143165 |
| ANT 1/2 | Sc-9299 Santa Cruz, RRID:AB_671086 |
| ANT2 | 14671 Cell Signaling, RRID:AB_2798562 |
| β-Actin | MAB1501 Millipore, RRID:AB_2223041 |
| β-3-Tubulin | sc-80005 Santa Cruz, RRID:AB_2210816 |
| Drp1 | sc271583 Santa Cruz, RRID:AB_10659110 |
| HKII | 2867, Cell Signaling, RRID:AB_2232946 |
| Mfn1 | sc-50330 Santa Cruz, RRID:AB_2250540 |
| OCT4 | 2840 Cell Signaling, RRID:AB_2167691 |
| Total OXPHOS Rodent Antibody Cocktail | ab110413 Abcam, RRID:AB_2629281 |
| PGC1-a | sc-13067 Santa Cruz, RRID:AB_2166218 |
| PKM2 | 4053 Cell Signaling, RRID:AB_1904096 |
| SDHA | ab14715 Abcam, RRID:AB_301433 |
| TFAM | sc-23588 Santa Cruz, RRID:AB_2303230 |
| TOM20 | sc-11415 Santa Cruz, RRID:AB_2207533 |
| TROMA-I | DSHB |
| Anti-Goat | sc-2354 Santa Cruz, RRID:AB_628490 |
| Anti-Mouse | 7076 Cell Signaling, RRID:AB_330924 |
| Anti-Rabbit | 7074 Cell Signaling, RRID:AB_2099233 |

### 2.5 Intracellular ATP Levels

Cells were seeded in white 96-well plates at a cellular density of 3.5×10^4^ cells/well, 24 hours before the experiment (24 hours after transfection). Four hours before the measurement, cells were incubated with media containing 2-DG at a final concentration of 50 nM, or with media as control, at 37°C, 5% CO_2_. Equal volume of Cell-Titer Glo Reagent (Cell-Titer Glo kit, Promega) was added, and luminescence was read after 2 minutes of orbital shaking and 10 minutes of incubation (in the dark) to allow luminescent signal stabilization, using Cytation3^TM^ UV–vis multi-well plate imaging reader.

### 2.6 Spheroid formation and migration assays

Upon 24h of transfection, cells were enzymatically harvested using trypsin and manually disaggregated to form a single-cell suspension. Three-dimensional cellular spheroids were prepared by seeding the cells in non-adherent conditions in 6 multi-well plates previously coated with Poly(2-hydroxyethyl methacrylate), at a density of 20,000 cells/well. Cells were allowed to grow for 1 day in High glucose Dulbecco’s modified Eagle’s medium (DMEM-D5648) supplemented with 7.5% FBS, 1.8 g/l sodium bicarbonate, 110 mg/l sodium pyruvate, and 1% antibiotic/antimycotic solution, at 37°C in a humidified atmosphere of 5% CO_2_. Spheroid-forming efficiency (SFE) was calculated as the ratio of formed spheroids above 60 μm relative to seeded cells. To assess the effect of etoposide on the Spheroid Forming Efficiency (SFE), transfected cells were seeded as described previously, with the addition of etoposide (0.25 μM). Subsequently, the formed spheroids were then collected and seeded in 96-well plates under adherent conditions, for 24 hours. The migration index was calculated as the ratio of the area at 24 hours to the area at 0 hours.

### 2.7 Cellular oxygen consumption and extracellular acidification measurements

Oxygen consumption rate (OCR) measurements and the extracellular acidification rate (ECAR) were measured using Seahorse XFe96 Extracellular Flux Analyzer (Agilent), using the Mito Stress and Glycolytic Rate protocols. Briefly, cell lines were seeded in Seahorse XF96 cell culture microplates previously coated with poly-D-lysine, at a final density of 35,000 cells/well, 24 hours before the experiment. On the day of the experiment, cells were washed with pre-warmed basal DMEM-D5030 (Sigma-Aldrich) medium supplemented with 25 mM glucose, 4 mM L-glutamine, 1 mM sodium pyruvate, and 5 mM HEPES (pH 7.4), and incubated at 37 °C in a non-CO_2_ incubator for 1 hour. OCR measurements were obtained in different respiratory states by sequential injection of 3 μM oligomycin, 0.25 μM FCCP, and 1 μM rotenone/antimycin A for the Mito Stress assay, and 1 μM rotenone/antimycin A, and 50 mM 2-DG the glycolytic rate assay. SRB was used to evaluate cell mass for normalization purposes. The results were analyzed using the Software Version Wave Desktop 2.6 and exported using the respective XF Report Generators (Agilent). The OCR and ECAR values were used to calculate the glycolytic parameters and expressed as Glycolytic Proton Efflux Rate (GlycoPER) and PER (Proton Efflux Rate), according to the manufacturer’s instructions.

For OCR and ECAR measurements on spheroids, spheroids were harvested after 24 hours of the spheroid formation assay, dissociated, and seeded under the same conditions as described above. After seeding the cells, the plate was centrifuged at 12000 rpm for 1 min to promote cell settling and then placed in the incubator at 37°C and 5% CO_2_ atmosphere for 1 hour before the experiment began.

The energy maps were also generated from the OCR and ECAR values under basal conditions and after induced stress following oligomycin and FCCP injection in 2D and 3D experiments. The raw data were also normalized to cell mass using the SRB method.

### 2.8 Measurement of mitochondrial membrane polarization

Transfected cells were seeded in μ-Slide 8 Well ibiTreat (80826, Ibidi) at a cellular density of 30,000 cells per well (24 hours after transfection). After 24 hours of seeding, cells were incubated with 100 nM Tetramethylrhodamine methyl ester (TMRM, Thermofisher) and 1 μg/ml Hoechst 33342 (Invitrogen, CA, USA), prepared in fresh complete growth medium, for 30 min, in the dark, at 37°C, and in a 5% CO_2_ atmosphere. Confocal images were collected at 63X magnification using a Plan-Apochromat lens (numerical aperture: 1.4), in Carl Zeiss LSM 710 laser scanning confocal microscope, pre-heated at 37°C, using Diode 405-30 (405nm) and DPSS 561-10 (561nm) lasers. Image processing and analysis were performed using ImageJ 1.51u software (RRID:SCR_003070).

### 2.9 Immunocytochemistry

P19 cells were seeded on glass coverslips at a cellular density of 30,000 cells (24 hours after transfection). Cells were then fixed with 4% paraformaldehyde in PBS (pre-warmed at 37LJC) for 15 min and permeabilized in 0.2% Triton X-100 in PBS for 2 min. Following a blocking step for 1 hour at room temperature with 3% BSA in PBS, cells were incubated with primary antibody against succinate dehydrogenase complex subunit A (SDHA) and HKII, diluted in blocking solution (1:100), overnight, at 4 °C. After 3 washes with PBS for 5 min, the secondary antibody Alexa Fluor-594 goat anti-mouse and Alexa Fluor-488 goat diluted in blocking solution (1:200) was applied for 1 hour at room temperature. Nuclei were stained with 1 μg/ml Hoechst 33342 (Invitrogen, CA, USA) in PBS for 10 min and coverslips were mounted using Dako (Sigma Chemical, St. Louis, MO, USA). Confocal images were acquired at 63X magnification using a Plan-Apochromat lens (numerical aperture: 1.4), in Carl Zeiss LSM 710 laser scanning confocal microscope. HKII expression was assessed using ImageJ 1.54h software (RRID:SCR_003070). Briefly, images underwent segmentation by thresholding the different channels. Binary images were generated to optimally resolve the mitochondria network and nucleus and further trace ROIs. Then, the mean gray value of HKII was measured within the ROIs, in the different cell compartments. SDHA labeling and the respective ROI were used to calculate the mitochondrial network area of each image, which was then normalized by the number of nuclei.

### 2.10 Statistical Analysis

All statistical analysis was performed using GraphPad Prism 8.0.2 software (RRID:SCR_002798). Data were checked for normal distributions and homogeneity of variances by the Shapiro-Wilk test and the F-test, respectively. Student’s t-test was used for statistical analysis of two means. In cases where normality was not verified, the Mann-Whitney test was performed. Two-way ANOVA with Sidak’s multiple comparisons was performed for statistical analysis to compare more than two groups with 2 independent variables when appropriate unless stated otherwise. Results are presented as means ± SEM. Significance was accepted with *p < 0.05, **p < 0.01, ***p < 0.0005, **** p < 0.0001.

## 3. Results

### ANT2 expression is upregulated in P19SCs

Despite advances in targeting CSCs, an effective and directed elimination of this cell population is still challenging. Using P19SCs, a previously described as a good reliable model to study CSCs and stemness features, Vega-Naredo et al., showed that total protein levels of ANT1/2 were differently expressed when compared to their differentiated counterparts [8]. However, the isoform responsible for that difference was not clarified. To address this, P19SCs were subjected to retinoic acid-induced differentiation (1 μM, 96 hours) and subsequently analyzed for pluripotency, differentiation markers, and morphological features. P19SCs exhibit small and round shape, reduced expression of trophectodermal cytokeratin 8 Endo-A (TROMA-I, marker for primitive endoderm) and β-3-Tubulin (marker for mature neurons), while the pluripotency marker octamer-binding transcription factor (OCT4) is overexpressed (Fig. 1A-C). After differentiation, the expression levels of the pluripotency marker OCT4, as well as of TROMA-I and β-3-Tubulin were highly increased in P19dCs, as expected. In addition, morphological changes were observed in P19dCs, with the cells exhibiting a more elongated shape (Fig. 1A-C). After the successful implementation of cell differentiation, ANT 1/2 and ANT2 protein levels were determined in both cell types. ANT 1/2 protein levels were strongly increased in P19dCs, corroborating previously described results (Suppl. Fig. 1). Nevertheless, the ANT2 isoform was found predominantly overexpressed in P19SCs, while expression was 75% lower in P19dCs (Fig. 1B-C), indicating that P19SCs have higher levels of ANT2 than their differentiated counterparts.

**Figure 1.**
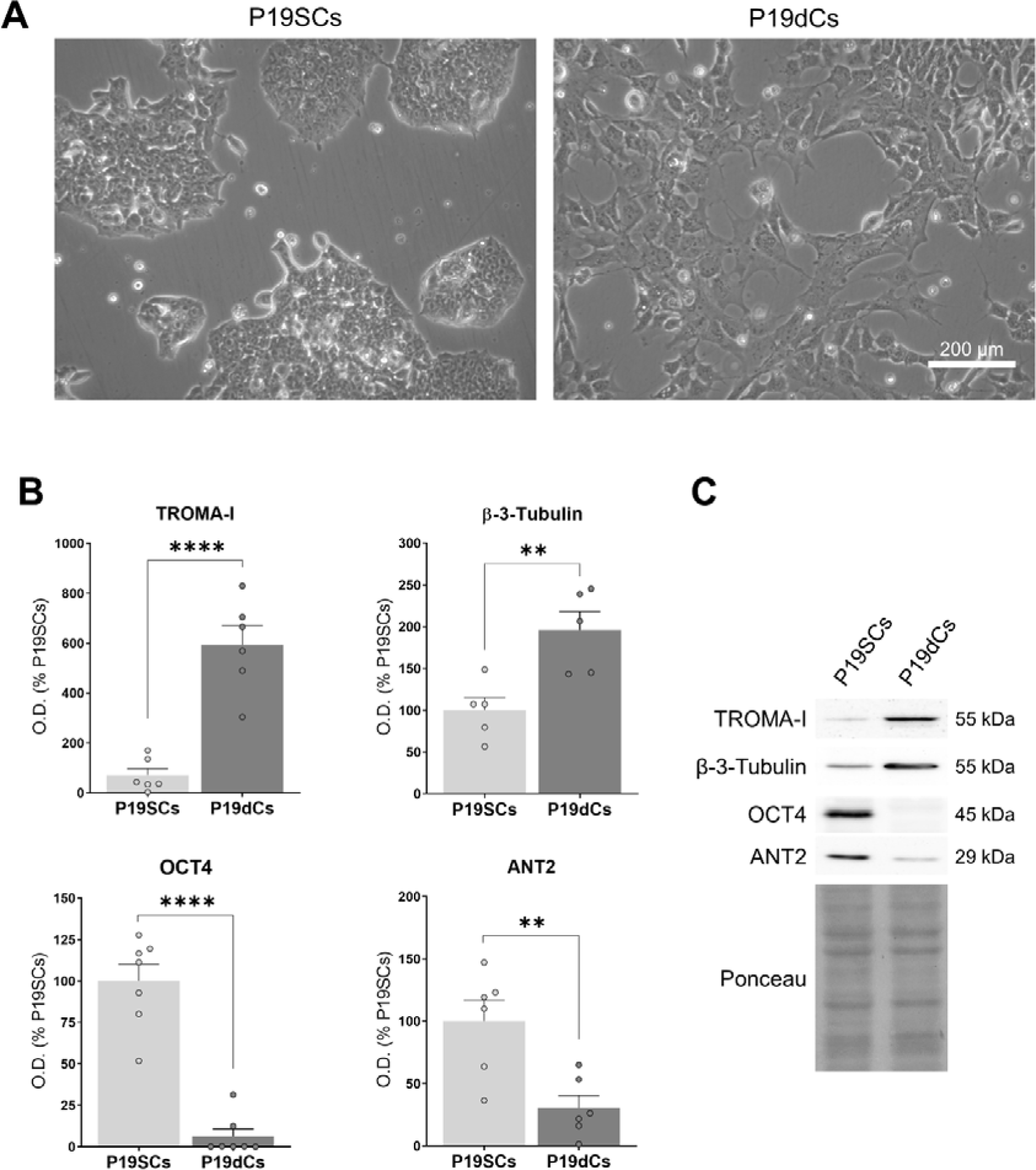
ANT2 expression levels in P19SCs and differentiated counterparts. A) Microscopy images of P19 embryonal carcinoma stem cells (P19SCs) and differentiated counterparts (P19dCs) upon 1 µM retinoic acid (RA) treatment for 96h. **B)** Differentiation (TROMA-I and β-3-Tubulin) and pluripotency (OCT4) markers, and ANT2 protein levels in P19SCs and P19dCs (1 µM, 96h), and **C)** representative Western blotting images. Ponceau S was used for loading control and normalization purposes. Data as mean ± SEM of optical density (O.D.) expressed as a percentage of P19SCs. Statistical analysis was performed using Student’s t-test. **p < 0.001, ****p < 0.0001 statistically significant to siCtrl.

### ANT2 protein expression levels affect P19SCs cell mass

Based on the above-described data, we wondered whether ANT2 could influence the P19SCs differentiation process. To this end, gene expression was transiently silenced with siRNA. Western blotting evidenced efficient protein knock-down up to 96h after transfection (Fig. 2A). To verify whether P19 cells maintained their stemness properties upon ANT2 suppression, the expression of pluripotency and differentiation markers as well as cell morphology were evaluated. The protein levels TROMA-I and OCT4 were not affected by ANT2 silencing (Fig. 2B). In terms of cell morphology, the transfected cells resembled P19SCs (Suppl. Fig. 2), which also suggests that the absence of ANT2 does not seem to compromise differentiation process of stem cells.

**Figure 2.**
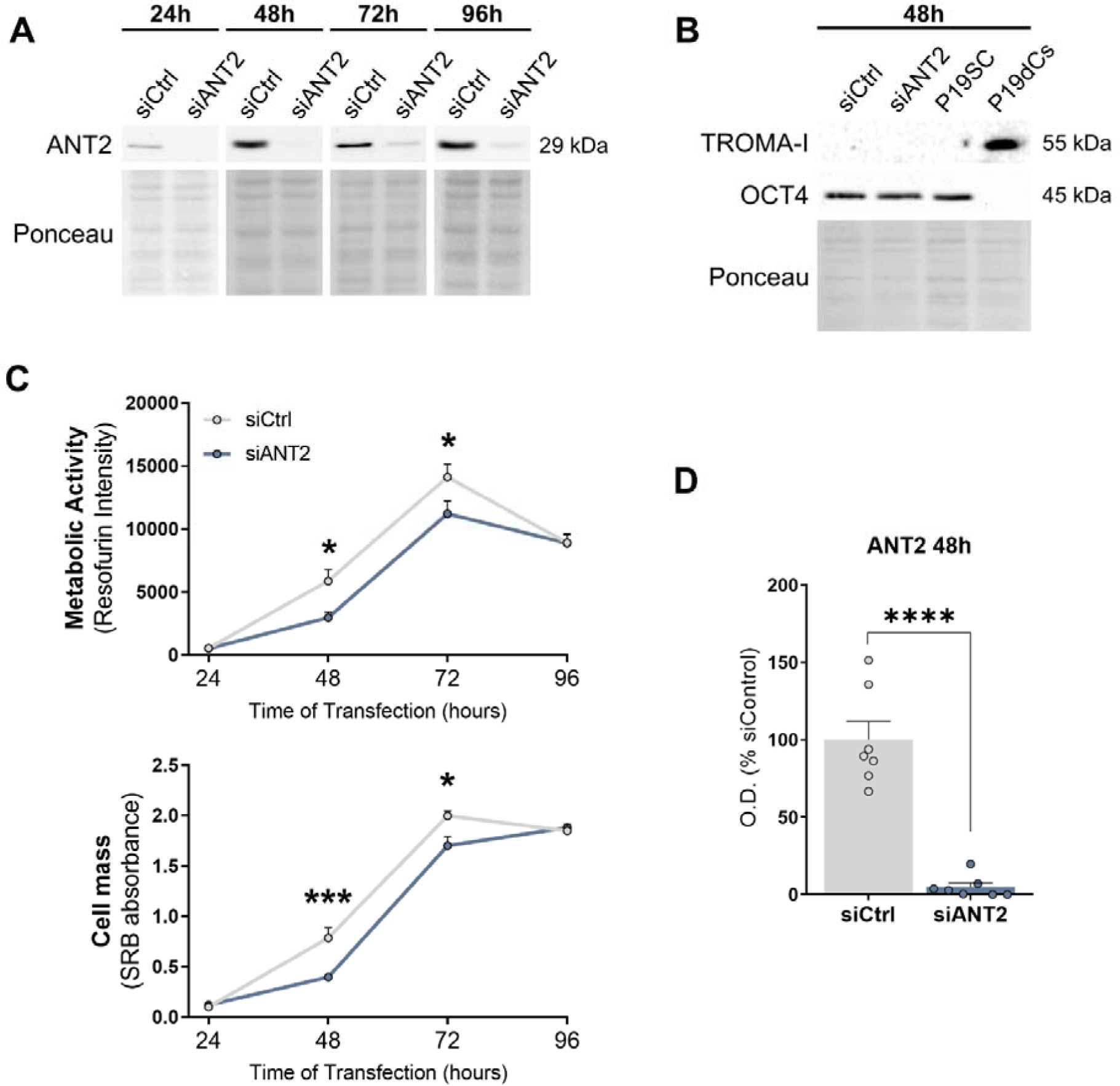
Characterization of P19 ANT2 silenced cells. A) Representative Western blotting image of ANT2 protein levels in P19SCs at several timepoints upon transfection. **B)** Differentiation and pluripotency markers after 48h of transfection. Ponceau S was used for loading control. **C)** Metabolic activity and cell mass measurements by resazurin and SRB after 24, 48, 72, and 96 h of transfection. Data as mean ± SEM, n ≥ 5 independent experiments. Statistical analysis was performed using a two-way ANOVA test with Sidak’s multiple comparisons. **D)** Expression levels of ANT2 after 48 h of transfection. Statistical analysis was performed using the Student’s test. Data as mean ± SEM. *p < 0.05, ***p < 0.005, ****p < 0.0001 statistically significant to siCtrl.

In addition to the generation of other cell types, CSCs are described as being fundamental for tumor growth, and several studies have shown that ANT2 is associated with a high proliferation rate [16,18], including in some tumor types [21,22]. The effect of ANT2 on the proliferation rate of transfected P19SCs was further evaluated using resazurin and SRB assays. A significant decrease in metabolic activity and cell mass was observed at 48 and 72 hours after ANT2 silencing on P19SCs (Fig. 2C). These results imply a critical role of ANT2 in the proliferation of cancer stem cell population. Importantly, the time point 48h seems to be the one with the most effective silencing (Fig. 2A and D), thus the following experiments were carried out after 48h ANT2 silencing.

### ANT2 silencing compromises P19SCs reliance on OXPHOS

ANT2 is increasingly expressed in undifferentiated cells and has been directly associated with glycolytic metabolism in cancer [13,19]. P19SCs are described as glycolytic and rely on this pathway for energy supply relative to their differentiated counterparts [8]. Glycolytic cancer cells import glycolysis-generated ATP into mitochondria via ANT2. Considering the impact of ANT2 on the proliferation of P19SCs, the cells’ metabolic profile was evaluated upon ANT2 gene downregulation. Glycolysis and mitochondria-related proteins were analyzed after 48 hours of ANT2 silencing. The expression of hexokinase II (HKII), the enzyme responsible for the first reaction of glycolysis, was decreased by 30% in siANT2 cells (Fig. 3A, B), while the pyruvate kinase muscle isoenzyme M2 (PKM2) levels, which catalyzes the last step, was unchanged (Fig. 3A, C). The observed differences in HKII protein expression prompted us to investigate ATP concentration in the P19SCs and siANT2 silencing cells on the glycolytic pathway. In basal conditions, there was no significant differences in the ATP levels between siCtrl and siANT2 cells (Fig. 3D). However, when 2-DG was used to inhibit glycolysis, a 38% depletion in ATP levels was observed in siCtrl and a 57% decrease in ANT2-silenced cells compared to vehicle-treated siCtrl cells. Notably, ATP levels were significantly lower in 2-DG-treated siANT2 cells compared to 2-DG siCtrl (Fig. 3D), indicating the importance of ANT2 in the energy metabolism of P19SCs.

**Figure 3.**
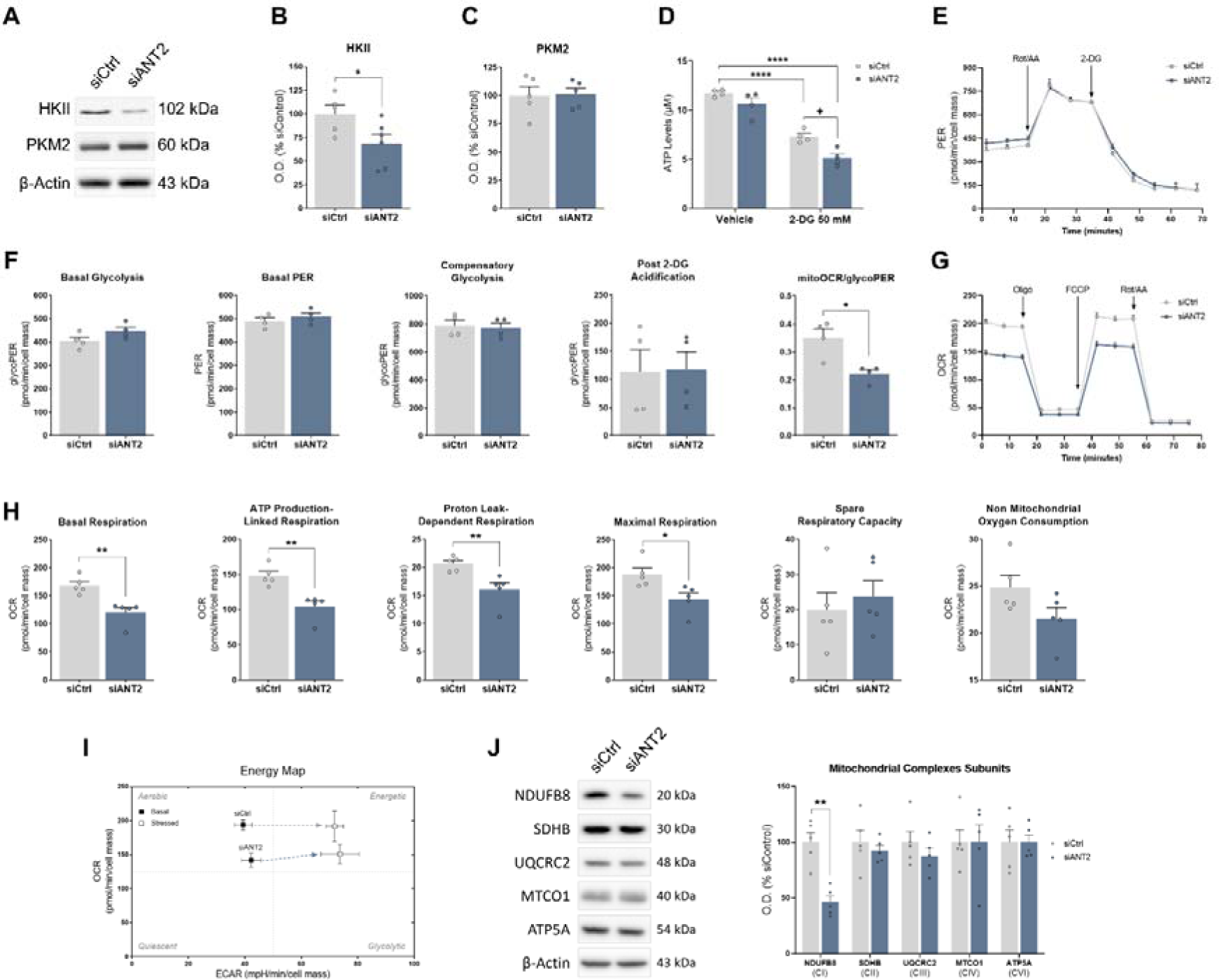
**P19SCs metabolic adaptations after ANT2 silencing**. A-C) HKII and PKM2 protein levels, and representative Western blotting images. Data as mean ± SEM of O.D. expressed as a percentage to siControl (siCtrl) after normalization with β-actin. D) Intracellular ATP levels in siCtrl and siANT2 cells after 4 h incubation with vehicle or 2DG. E) Representative image of proton efflux rate (PER) measurement in siCtrl and siANT2 cells (48 h post-transfection), with injection of rotenone + antimycin (Rot/AA, 1 µM) and 2-DG (50 mM). F) Basal glycolysis, basal proton efflux rate, compensatory glycolysis, post-2-DG acidification, and mitoOCR/glycoPER rate measurements. F) Representative image of oxygen consumption rate (OCR) measurement in siCtrl and siANT2 cells (48 h post-transfection), with injection of oligomycin (oligo, 3 µM), FCCP (0.25 µM), and Rot/AA (1 µM). H) Basal respiration, ATP production-linked respiration, proton leak-dependent respiration, maximal respiration, spare respiratory capacity, and non-mitochondrial oxygen consumption of P19Scs 48 h after transfection. Data from Seahorse analysis as mean ± SEM. n ≥ 4 independent experiments (4-6 replicates each). I) Energy map obtained by plotting the OCR and ECAR measurements under basal and stressed (upon oligomycin and FCCP exposure) conditions. J) Representative images of Western blotting and protein expression levels of mitochondrial complexes subunits 48h after ANT2 silencing. Data as mean ± SEM expressed as a percentage to siCtrl. Statistical analysis was performed using the Student’s test to compare two mean values and the two-way ANOVA with Sidak’s multiple comparison post-test to compare more than two groups with 2 independent variables. *p < 0.05, **p < 0.01 and ****p < 0.0001 statistically significant to siCtrl. + p < 0.05 statistically significant to siCtrl 2-DG treated group.

To gain better insights into the glycolytic pathway, the metabolic activity of P19SCs was analyzed 48 hours after transfection using the Seahorse XFe96 Extracellular Flux Analyzer, as the effects on cell proliferation and intracellular ATP content were most pronounced. Using the glycolytic rate assay, the proton efflux rate (PER) attributable to glycolysis (glycoPER) was determined by the extracellular acidification rate (ECAR) and oxygen consumption rate (OCR) measurements. Interestingly, no differences were observed in glycolysis with ANT2 suppression (Fig. 3E, F, I). However, the mitoOCR/glycoPER ratio was decreased in siANT2 cells, which prompted us to ask whether ANT2 depletion could interfere with mitochondrial respiration. To answer this question, the Mitostress assay was performed. Although no significant changes in non-mitochondrial oxygen consumption and spare respiratory capacity were detected, basal respiration, proton leak-dependent respiration, maximal respiration, and ATP production-linked respiration were significantly decreased in ANT2-silenced cells (Fig. 3G,H). The replotting of data from unstressed and stressed cells into an energetic map allowed to confirm the ANT2 effect on OCR and glycolytic profile (Fig. 3 I). Considering the above data, we next analyzed the expression of the mitochondrial electron transport chain subunits to verify possible changes in the OXPHOS machinery. The expression levels of NADH dehydrogenase [ubiquinone] 1 β subcomplex subunit 8 (NDUFB8), succinate dehydrogenase [ubiquinone] iron-sulfur subunit B (SDHB), cytochrome b-c1 complex subunit 2 (UQCRC2), cytochrome c oxidase subunit I (MTCO1) and ATP synthase subunit α (ATP5A), respectively from Complex I to V, were analyzed by Western blotting 48 hours after transfection. The protein levels of Complex I subunit NDUFBB8 were reduced by 50% in ANT2-silenced P19SCs (Fig. 3J), in contrast to the other complexes, which showed no alterations. Overall, the data suggest that ANT2 levels influence the total HKII content and the composition of the mitochondrial respiratory machinery by promoting a change in mitochondrial respiration composition that may affect cells’ fate.

### ANT2 silencing induced changes in Hexokinase II protein levels

Hexokinases are primarily associated with glucose metabolism, with overexpression of HKII being particularly associated with cancer [25]. However, the multifaceted role of HKII extends beyond this, encompassing its role as a mitochondria-docked protein, initiation of mitochondria-mediated apoptotic cell death, coexpression with ANT2, and involvement in cancer stemness through nuclear localization and regulation of calcium fluxes [26–28]. Therefore, clarification of HKII expression and subcellular localization in P19SCs after silencing ANT2 was considered of utmost importance.

Although we found no differences in most glycolytic-related parameters (Fig. 3), a significant decrease in HKII protein expression and ATP levels was observed in siANT2 cells (Fig. 3A, B). Using confocal imaging, we confirmed a decrease in the total amount of HKII in siANT2 cells, consistent with the previous data from Western blot analysis (Fig. 4A, B). Moreover, the subcellular localization of HKII labeling evidenced that under ANT2 silencing, HKII levels were decreased in both nucleus and mitochondria, as well as outside both organelles (Fig. 4A,B). This data shows that ANT2 in the P19 cell line causes widespread regulatory repression of HKII expression in the cell.

**Figure 4.**
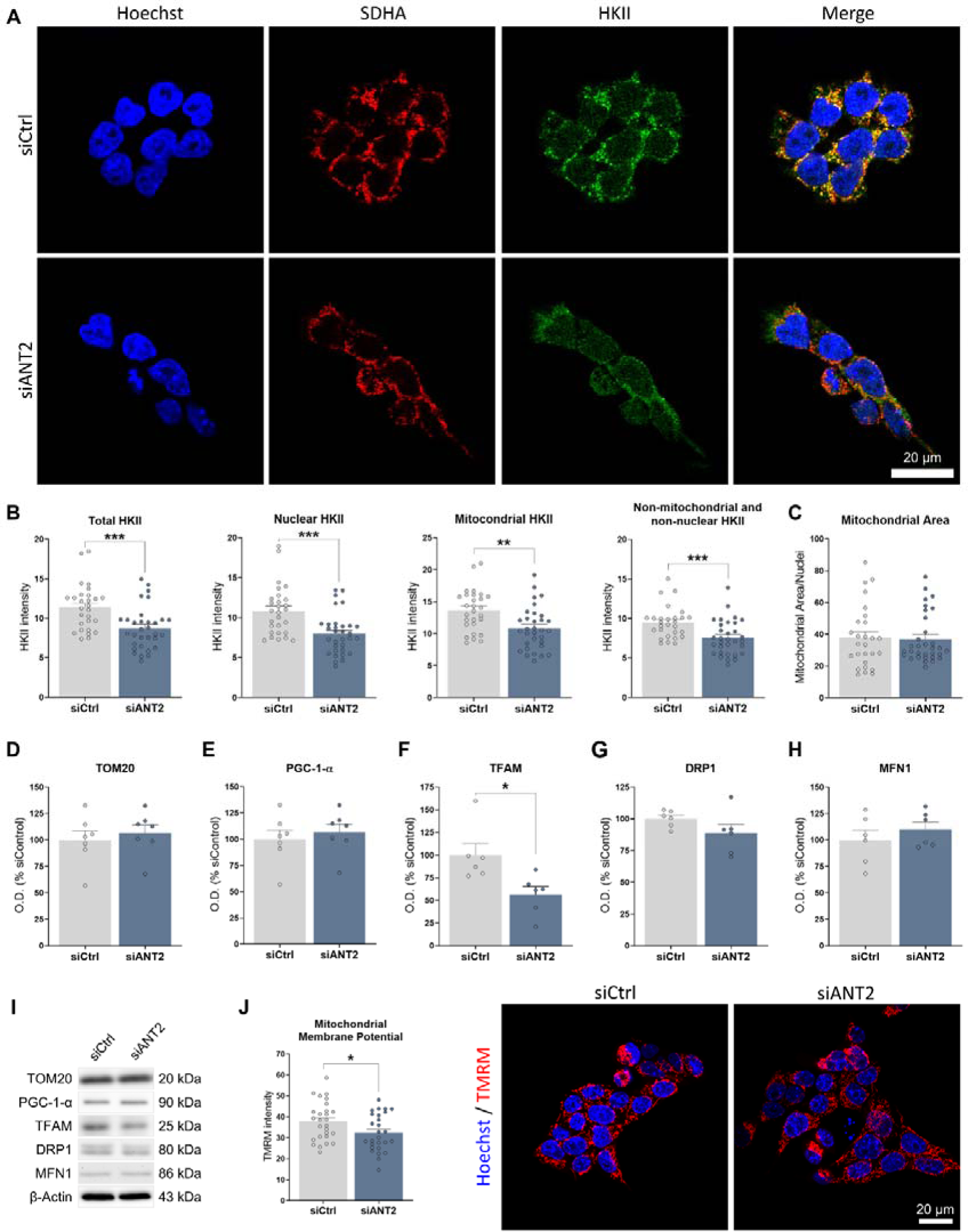
**ANT2 silencing effect on HKII subcellular localization and mitochondrial parameters**. A) Representative confocal microscopy images of SDHA (mitochondrial marker), HKII, and nucleus stain (Hoechst 33342) in P19SCs transfected cells, from 3 independent experiments. B) HKII fluorescence intensity analyzed across the entire cell, in the nucleus, in the mitochondria, and in non-mitochondrial and non-nuclear localization. C) Mitochondrial network area measured by the area of SDHA labeling. D) TOM20, E) PGC-1-α, F) TFAM, G) DRP1, and H) MFN1 protein levels quantification, expressed as percentages to siCtrl, and I) the representative Western blotting images. J) Mitochondrial membrane potential measured by TMRM fluorescence intensity, and the representative confocal microscopy images, from 6 independent experiments. Data as mean ± SEM. Statistical analysis was performed using the Student’s test. *p < 0.0, **p < 0.01, ***p < 0.0005 statistically significant to siCtrl.

### Switching off ANT2 lowers the mitochondrial potential without affecting the mitochondrial network area

To gain a more comprehensive understanding of the mitochondrial network after siANT2 silencing, markers of mitochondrial mass, biogenesis, and dynamics were analyzed. The mitochondrial area per cell was quantified by labeling SDHA as a mitochondrial marker (Fig. 4A, C). No significant differences were observed between siANT2 and siCtrl cells, consistent with the protein levels data of translocase of outer mitochondrial membrane 20 (TOM20) (Fig. 4D, I). While protein levels of peroxisome proliferator-activated receptor-gamma coactivator 1-alpha (PGC-1-α) remained unchanged, a 50% decrease in mitochondrial transcription factor A (TFAM) protein expression was observed in the absence of ANT2 (Fig. 4E, F, I). Since the mitochondrial network is related to fusion and fission mechanisms and has been described as an important component of tumor initiation and progression [29–31], we further evaluated it by Western blotting. No differences were found for mitofusin 1 (MFN1), a fusion protein, nor for dynamin-related protein-1 (DRP1), a fission protein (Fig. 4G-I). These findings suggest that ANT2 depletion does not disrupt mitochondrial mass or dynamics. However, it appears to influence mitochondrial biogenesis, at least in the short term, as indicated by the decreased levels of TFAM.

It is hypothesized that ANT2 is responsible for ATP translocation to the mitochondria under pathological conditions, such as in cancer [32]. Therefore, ATP uptake into mitochondria could be impaired after ANT2 silencing, which could affect the maintenance of the mitochondrial membrane potential in P19SCs. Using the TMRM probe, which accumulates in negatively charged polarized mitochondria [33], the mitochondrial polarization was evaluated by confocal microscopy. As expected, ANT2 silenced cells exhibited a decreased mitochondrial polarization, as shown by the TMRM fluorescence intensity and representative images (Fig. 4J). These findings suggest that ANT2 silencing may impair ATP uptake and subsequent disruption of the mitochondrial membrane potential, highlighting the potential role of ANT2 in maintaining mitochondrial function under pathological conditions.

### ANT2 depletion impairs stem cell spheroids’ efficiency formation and migration

CSCs possess the ability to form spheroids when cultivated under non-adherent conditions. This reflects their self-renew capacity and clonogenic potential, which are fundamental characteristics for tumor initiation, progression, and therapeutic resistance [34]. Furthermore, by employing a 3D model, we can better recapitulate CSCs and solid tumors’ biological characteristics *in vitro*. To assess the impact of ANT2 depletion in 3D model tumor formation, P19SCs were cultured under non-adherent plates coated with poly-HEMA to assess spheroid-forming efficiency. Interestingly, upon ANT2 depletion confirmation in the 3D approach (Fig. 5A), we observed a 30% reduction in spheroid formation efficiency in ANT2 silenced cells (Fig. 5B). This finding highlights the potential involvement of ANT2 in regulating CSC behavior and tumor growth in 3D structures.

**Figure 5.**
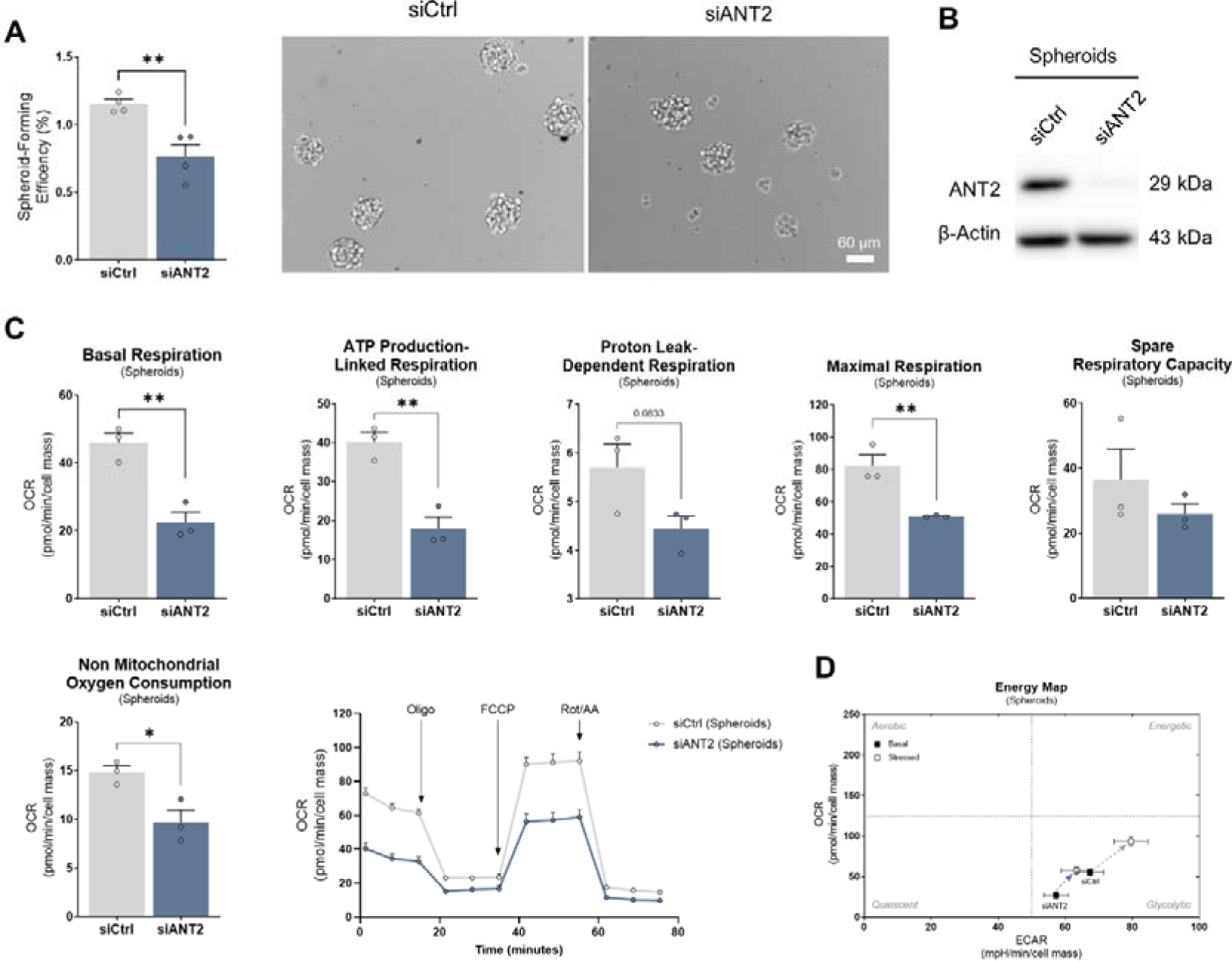
Effect of siANT2 on spheroids formation and mitochondrial respiration. A) Spheroid-forming efficiency of siCtrl and siANT2 cells, and the respective representative pictures of the spheroids. B) Representative Western blotting of ANT2 protein levels in the spheroids formed from siCtrl and siANT2 cells. C) Basal respiration, ATP production-linked respiration, proton leak-dependent respiration, maximal respiration, spare respiratory capacity, and non-mitochondrial oxygen consumption of siCtrl and ANT2 cells from the spheroids, and the representative kinetics graph of oxygen consumption rate (OCR) measurement in siCtrl and siANT2 cells, with sequential addition of oligomycin (oligo, 3 µM), FCCP (0.25 µM), and rotenone + antimycin (Rot/AA, 1 µM). D) Energy map obtained by plotting the OCR and ECAR measurements under basal and stressed (upon oligomycin and FCCP exposure) conditions. Data are presented as mean ± SEM of 3-4 replicates, from 3 independent experiments. Statistical analysis was performed using the Student’s test. * p < 0.05 and ** p < 0.01 statistically significant differences.

We further investigated ANT2 depletion effects in the metabolic status of siCtrl and siANT2 spheroids, similarly to monolayer analysis. Remarkably, spheroids showed a similar trend on OCR and GlycoPER parameters with no significant differences in the glycolytic rate but a decreased reliance on OXPHOS (Fig. 5C, Supp. Fig 3).. Notably, the trend was even more pronounced in ANT2-silenced spheroids when compared to monolayer culture. Specifically, ANT2-silenced spheroids presented a 50% decrease in OCR relative to siCtrl, whereas only a 30% reduction was observed in the monolayer. Interestingly, spheroids’ energy map accompanied these results although, under stress conditions, ANT2 expression seems to promote further P19SCs reliance on glycolytic metabolism while OCR differences are less visible (Fig. 5C, D, Supp. Fig 3). From these data, it appears that the expression of ANT2 in 3D cultures strongly influences the metabolism of CSCs, which underscores the gene effect in stem cells’ ability to form spheroid and thus self-renewal and metastasis potential.

To explore the potential synergistic effect of ANT2 silencing and chemotherapeutic drugs on CSCs and their ability to form spheroids, cells were also cultured with etoposide (0.25 μM) under non-adherent conditions. Interestingly, etoposide alone significantly reduced the efficiency of spheroid formation in the siCtrl group by 45% (Fig. 6A). Furthermore, the combined effect of siANT2 plus etoposide treatment induced a 30% decrease in stem cells’ ability to form spheroids, in comparison to only etoposide-treated cells (Fig. 6A). Afterwards, the collected spheroids were cultivated under adherent conditions to assess their migratory capacity. Consistently, we observed a significant decrease in the migration index of spheroids treated with siANT2 or etoposide (approximately 15% and 45%, respectively), in comparison to vehicle-treated spheroids (siCtrl vehicle) (Fig. 6B, C). Moreover, spheroids treated with siANT2 plus etoposide also exhibited a statistically significant decreased migration compared to etoposide-treated siCtrl spheroids (20%, Fig. 6B, C). It is also important to mention that at 24 hours after the seeding of spheroids treated with etoposide, many apoptotic bodies were seen, both in siCtrl and siANT2 spheroids. These results suggest that etoposide inhibits the spheroid formation and the migratory ability of CSCs, possibly due to its cytotoxic properties targeting rapidly dividing cells, which is further enhanced by ANT2 silencing.

**Figure 6.**
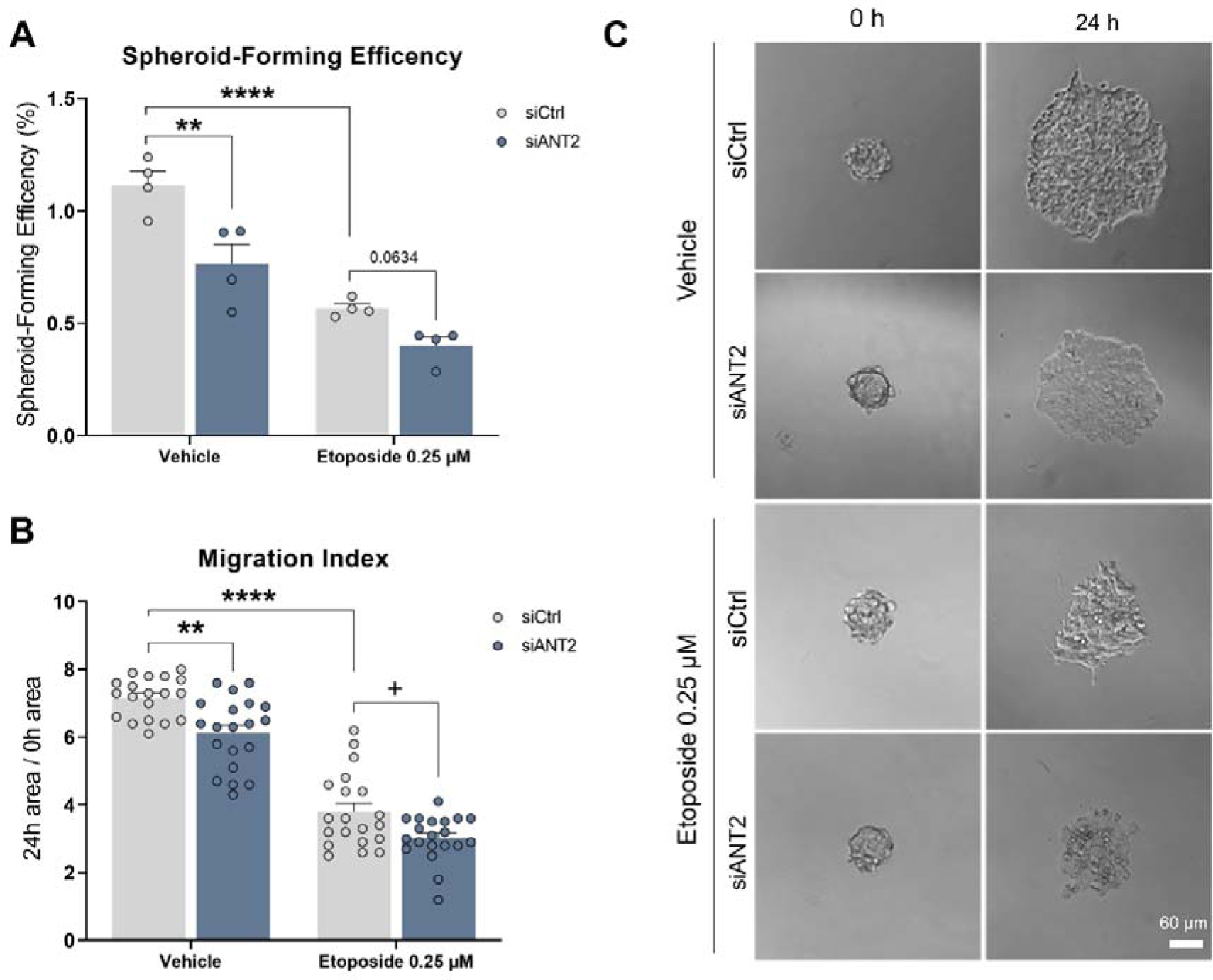
Synergetic effect of ANT2 silencing and etoposide on P19SCs sphere-forming efficiency and migration. A) Spheroid-forming efficiency of siCtrl and siANT2 cells in the presence of etoposide (0.25 μM, 24 h). B) Migration index of the spheroids upon seeding on adherent conditions. C) Representative images of the spheroids under adherence conditions (0 and 24 h). Data as mean ± SEM. Statistical analysis was performed using the two-way ANOVA with Holm-Sidak’s multiple comparison post-test to compare more than two groups with 2 independent variables. ** p < 0.01 and **** p < 0.0001 statistically significant to the siCtrl vehicle group. + p < 0.05 statistically significant to siCtrl-etoposide treated group.

## 4. Discussion

Endowed with unique self-renewal and differentiation abilities, CSCs are regarded as persistent factors supporting tumor heterogeneity, resistance, and metastatic potential [35,36]. Despite the relevant advances and accumulated knowledge in recent years regarding their biological characteristics such as cell surface biomarkers, signaling pathways, or metabolism, much is still unknown. Thus, a better understanding of the factors driving cancer growth, cell death resistance, metabolic remodeling, and high metastatic behavior is essential to effectively target tumor onset, progression, and response to treatments [36–38].

Using P19SCs as a convenient model to gain insights into CSCs and ANT2 function [9,39], we successfully validated overexpression of ANT2 isoform protein in P19SCs, contrasting with a decrease observed upon their differentiation, which is consistent with Vega-Naredo et al previous findings at the mRNA level [8]. These results align with studies demonstrating ANT2 upregulation in proliferative [18,40], undifferentiated [41], and cancer cells [13,42], such as lung, liver, neuroblastoma, prostate, or breast [19,21,43–47], including in cancer stem cells [22]. Thus, not surprisingly, it has been pointed out as a predictive prognosis biomarker in several tumor types [20,44,46,48].

Following our findings and the perspective that tumor growth arises from the persistent differentiation capacity of CSCs to generate progeny, we show that silencing ANT2 in P19 cells had a suppressive effect on their proliferation, but without affecting stem cell morphology and pluripotency/differentiation markers. ANT2 siRNA induced a significant decrease in proliferation rate, especially after 48 hours of transfection, reaching a 50% decrease compared to siCtrl cells, supporting its role in CSCs cell proliferation. Recent studies also demonstrate that ANT2 depletion impairs tumor growth of breast [21] and prostate cancer [47], both *in vitro* and *in vivo*. This is consistent with ANT2’s established involvement in cell growth, and its recognition as a proliferation biomarker [18].

ANT2 plays an important role in the metabolic adaptation of cancer cells during tumorigenesis, being correlated with the aggressiveness of cancer cells [8]. Despite ANT2’s association with higher glycolytic phenotype [13,49] and ATP depletion in breast cancer cells [21], our data on glycolysis parameters on the highly glycolytic P19SCs remained unchanged by ANT2 silencing, in both 2D and 3D conditions. ATP levels were significantly decreased upon treatment with the glucose analog 2-DG, being further stimulated when combined with siANT2 treatment. This suggests that P19SC ANT2-depleted cells remain glycolytic dependent, while ANT2 may positively prompt an energetic decay if coupled with drugs targeting glycolysis. Although the total HK2 protein level was significantly affected by ANT2 silencing, the activity of this essential kinase for the glycolytic pathway was not measured and the PKM2 levels remained unchanged.

Interestingly, we demonstrated that ANT2 silencing significantly diminished mitochondrial respiration parameters and membrane potential in P19SCs. The ANT2 depletion and complex I subunit protein decreased expression likely interfere with the proton gradient generation across the inner mitochondria, thus explaining the decreased mitochondrial polarization and impairment of OXPHOS on the P19SCs[50]. The NDUFB8 subunit is required for Complex I full assembly [51], and mutations can lead to respiratory chain complex I deficiency [52]. Downregulation of this subunit, as seen in gastric adenocarcinoma cells exposed to γ-Tocotrienol, also prompts OXPHOS inhibition [53]. Evidence also suggests that HKII binding to VDAC works reversely, using cytosolic glucose-6-phosphate to produce ATP, which is then uptake into mitochondria by ANT2 to be used to repolarize mitochondria [13,54]. These studies sustain the lower OXPHOS function on siANT2 cells. Additionally, cancer cells commonly exhibit higher mitochondrial potential, which is associated with tumorigenic potential [55,56]. Notably, previous studies have shown that ANT2 silencing suppresses the mitochondrial potential of breast cancer cells [21], and is associated with apoptosis induction [67].

Maintenance of mitochondrial homeostasis results from the coordinated interplay of two opposing processes: mitochondrial biogenesis, which generates new mitochondria; and mitophagy, which eliminates damaged mitochondria. Several factors such as metabolic state, tissue type, and tumor heterogeneity, regulate this process [57]. Mitochondrial biogenesis is driven by PGC-1-α, which activates different transcription factors that ultimately promote the expression of TFAM, which is responsible for mtDNA transcription, replication, and maintenance [58,59]. TFAM levels were decreased in ANT2-silenced cells, but mitochondrial network area, TOM20, and PGC-1-α protein levels remained unchanged. Abnormal transcription of mitochondrial transcription factors and OXPHOs components leads to metabolic reprogramming, which is related to carcinogenesis. Despite controversial reports on TFAM effects on tumor progression [60], TFAM depletion elicits metabolic reprogramming in head and neck cancer and hepatocellular carcinoma (HCC) by impairing mitochondrial respiration [77,78]. In HCC, decreased TFAM content is allied with chemosensitization, and growth arrest without compromising glycolysis [78]. Thus, our findings support a view where ANT2-induced metabolic reprogramming in P19SCs may also be related to the modulation of TFAM content, rather than exerting a modulatory effect on mitochondrial biogenesis.

Another crucial aspect of CSCs is their ability to survive under non-adherent conditions due to anoikis resistance, thereby influencing tumor cells’ invasiveness capacity and propelling (re)growth [49]. The Bcl2 protein family is known to modulate the activity of ANT isoforms, including ANT2. Interestingly, ANT2 silencing has been found to trigger apoptosis in breast cancer and breast cancer-stem-like cells [21,22], suggesting ANT2 as an anti-apoptotic protein. Despite its role in glycolysis, HKII also binds to mitochondria by directly interacting with VDAC to stabilize a closed mPTP conformation [50], and its dissociation from mitochondria is reported to cause cell death by mitochondrial-induced apoptosis [51,52]. Jang et. al demonstrated that ANT2 depletion changes the balance of the Bcl-2 family in the mitochondrial membranes of breast cancer cells, leading to cell death by favoring a proapoptotic pore-forming state [42]. Since HKII and Bcl-2 protein family are considered part of the mPTP, both can be altered and play a role in mitochondria-initiated apoptotic cell death [53,54]. Here, we showed that ANT2 depletion induced an overall decrease in HKII protein content, including in mitochondria and nucleus. While previous studies have proposed ANT2’s indirect regulation of HKII transcription [55], further studies are required to validate this hypothesis, although our findings seem to reinforce it. The HKII mitochondrial detachment suggests an ANT2-dependent involvement of HKII in mitochondria-initiated apoptotic cell death of P19SCs. The association of HKII with the resistance of cancer stem-like cells to anoikis is reported in intrahepatic cholangiocarcinoma [56]. The nuclear localization of HKII in P19SCS, observed by us, is well documented in yeast and mammal cells, including cancer cells [57–61]. Recently, Thomas et al. showed that leukemic hematopoietic stem cells express high nuclear levels of HKII, being involved in the regulation of the accessibility to chromatin and reduction of double-stranded DNA breaks occurrence, which was suggested as a potential explanation for their enhanced chemoresistance [62]. Thus, nucleus-decreased HKII levels in ANT2-depleted P19SCs may represent an additional explanation for ANT2 effects in P19SCs.

By evaluating P19SCs’ ability to form spheroids, we demonstrated that ANT2 is an essential protein for CSCs’ survival in harsh environments and drug sensibilization. ANT2 knockdown led to a strong suppression of spheroid efficiency formation in P19SCs, in agreement with the previous findings reported in breast cancer stem cell subpopulation [22]. Etoposide promoted a more prominent decrease in spheroid formation than siANT2 alone, and while their combined effects were not significant, a positive synergistic effect (20% decrease) was observed. Accordingly, P19SCs ANT2-silencing and drug combined experimental approach particularly affected cell migration capacity, highlighting ANT2 as a potential therapeutic target. Recently, Zhang et al showed that genetic knockdown of ANT2 effectively inhibited proliferation, migration, and invasion of prostate cancer cells [47]. Etoposide is a commonly used chemotherapeutic drug for the treatment of different cancers, including stem cells, with resistance being also reported [38,63]. Combinatory approaches represent a promising strategy to overcome CSCs resistance to therapy [64]. Thus, our data highlight ANT2 as a relevant target for stem cells in tumor development and acquired resistance phenotypes, in agreement with evidence from non-small lung cancer and breast cancer studies [22,44].

Our findings in P19SCs denote the critical role of ANT2 in fundamental cellular events and stem cell features while shedding light on the molecular mechanisms that trigger these effects. This study shows that silencing ANT2 promotes metabolic reprogramming in P19SCs by modulating the OXPHOS machinery and mitochondrial membrane potential. These alterations likely occurred due to the changes d in HKII, complex I, and TFAM levels, resulting in cell growth arrest, anoikis, and drug sensitization of P19SCs. This suggests that targeting ANT2 and exploiting its associated vulnerabilities may offer a promising approach for controlling CSCs and managing tumor progression.

## Author contribution

**Gabriela L. Oliveira**: Methodology, Formal analysis, Investigation, Writing-Original draft; **Sandra I. Mota**: Formal analysis, Investigation, Writing-Review Editing; **Paulo J. Oliveira**: Conceptualization, Writing-Review and Editing, Supervision, Funding; **Ricardo Marques**: Conceptualization, Writing-Original draft, Writing-Review and Editing, Supervision, Visualization.

## Acknowledgments

Work in the authors’ laboratory is funded by FEDER—Fundo Europeu de Desenvolvimento Regional through the COMPETE 2020—Operational Programme for Competitiveness and Internationalisation (POCI), Portugal 2020, and by Fundação para a Ciência e Tecnologia (FCT), under the project POCI-01-0145-FEDER-016390 [CANCEL STEM], and through the COMPETE 2020 - Operational Programme for Competitiveness and Internationalisation and Portuguese national funds via FCT – Fundação para a Ciência e a Tecnologia, under project[s] UIDB/04539/2020, UIDP/04539/2020 and LA/P/0058/2020. SIM is supported by FCT (Ref: DL57/2016/CP1448/CT0027) and GLO is supported by FCT (Ref: 2020.04765.BD)

## Data availability

The datasets generated in this work are available at the figshare repository (https://figshare.com/account/home%23/projects/208600)

## Supplementary material

**Suppl. Fig 1.**
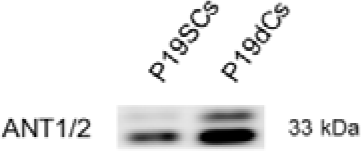
Representative Western blotting image for ANT1/2 P19SCs and P19dCs.

**Suppl. Fig 2.**
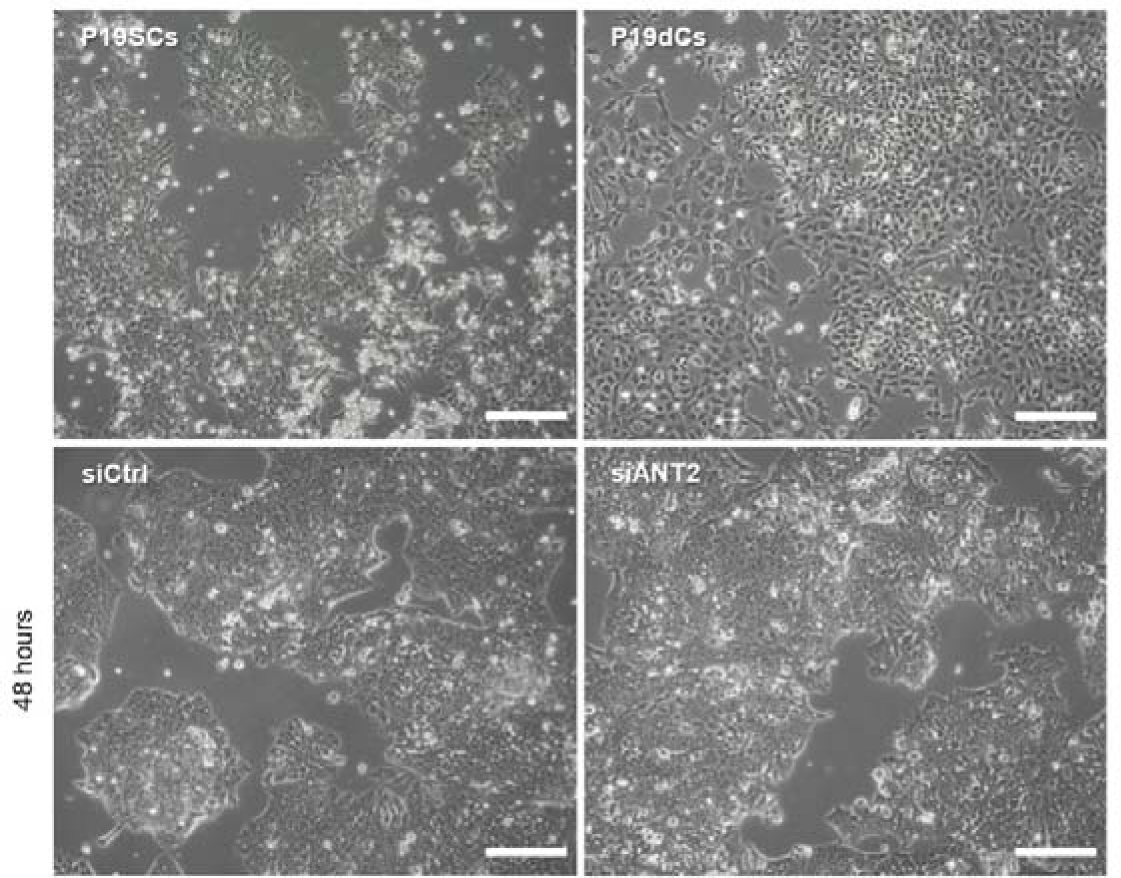
Microscopy images of P19SCs, P19dCs, siCtrl, and siANt2 cells, for comparison of morphology features. Scale bar: 200 μM.

**Suppl. Fig 3.**
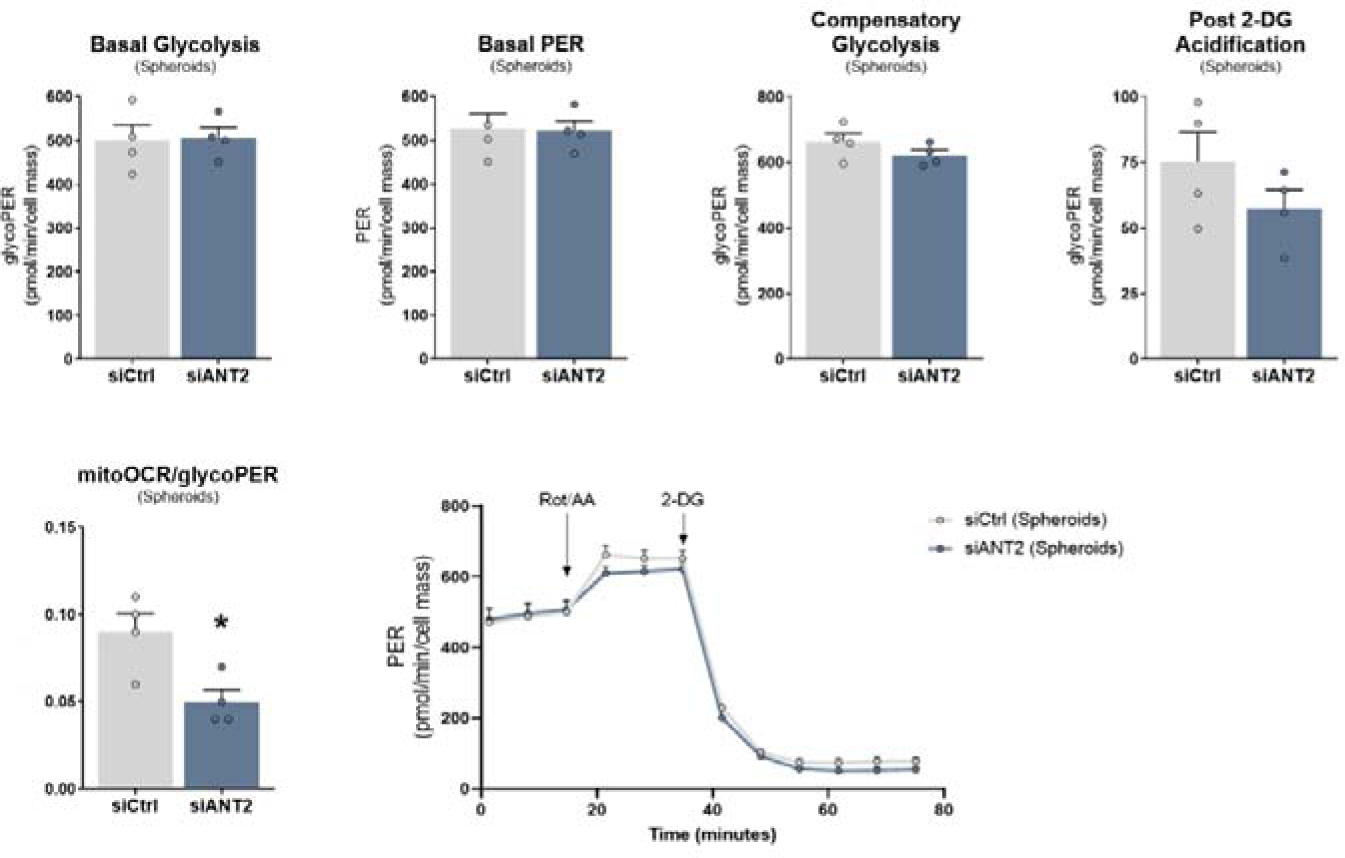
Basal glycolysis, basal proton efflux rate, compensatory glycolysis, post-2-DG acidification and mitoOCR/glycoPER rate measurements, and the respective image of proton efflux rate (PER) measurement in siCtrl and siANT2 spheroids, with injection of rotenone + antimycin (Rot/AA, 1 µM) and 2-DG (50 mM).

